# An auditory “low road” for threat in humans sensitive to fast temporal cues

**DOI:** 10.64898/2025.12.24.690990

**Authors:** Martina T. Cinca-Tomás, Emmanouela Kosteletou-Kassotaki, Jordi Costa-Faidella, Carles Escera, Judith Domínguez-Borràs

## Abstract

Rapid threat processing is crucial for survival. In vision, neural models of emotion propose the existence of direct pathways, or “low roads”, from visual thalamus to the amygdala, which may rely on magnocellular inputs and thereby enable rapid processing of threat signals. In audition, evidence from non-human animals points to the existence of a similar fast direct pathway, but its presence in humans remains unknown. Building on animal evidence that high-rate amplitude-modulated (highAM) sounds preferentially engage magnocellular processing, we used behavioral, physiological, and neuroimaging measures to search for an auditory “low road” for threat in humans responsive to magnocellular-associated features. Specifically, we tested whether highAM threat cues elicit rapid affective responses, and whether highAM threat processing aligns with structural thalamo-amygdala connectivity. We recorded electroencephalography and pupillometry from participants while they detected voices during fear conditioning. Emotional voices presented at highAM rates, as opposed to lowAM, elicited threat-related responses during the earliest post-stimulus window. A further study combining functional and diffusion-weighted imaging revealed that selective amygdala responses to highAM threatening voices were functionally associated with individual fiber density of an auditory thalamo-amygdala pathway. These results provide convergent functional and anatomical evidence supporting a subcortical pathway for auditory threat processing, consistent with evolutionarily conserved emotional systems.

## Introduction

Rapid threat processing is one of the most essential and evolutionarily adaptive functions of the human brain. In vision, one of the most extensively studied neural pathways serving this function is the so-called “low road” for emotion processing^1,2^. This phylogenetically ancient route ^3^ consists of direct anatomical projections from visual thalamic nuclei to the amygdala (a key structure for detecting danger)^4–7^, and allows rapid and coarse evaluation of threat signals before detailed cortical analysis^1,2,8–10^. In humans, this neural “shortcut” is thought to rely primarily on magnocellular visual inputs, which are strongly associated with low-spatial-frequency (LSF) information conveying coarse stimulus features^8,11–13^. Consistent with fast magnocellular transmission, human electrophysiology has revealed short-latency amygdala and cortical responses during LSF emotional face processing^13–15^ (see reviews in refs. ^8,9^). This route is believed to mediate abundant fear-associated physiological processes and behaviors^2,8,10,16^. Notably, patients with cortical blindness due to bilateral occipital damage retain unconscious fear processing presumably through these preserved subcortical routes^9,16,17^, showing amygdala responses predominantly for LSF fearful faces^18,19^. Altogether, broad evidence supports a magnocellular-sensitive visual “low road” that provides a rapid warning system for threat, a notion that has influenced affective neuroscience for several decades.

An intriguing question that remains unresolved is whether a comparable fast route for fear processing also exists in human audition. In non-human species^1,20–23^, extensive anatomical and functional evidence points to an auditory “low road”^24^, with direct projections from the medial geniculate body (MGB, the auditory thalamus) to the basolateral amygdala (BLA), a pathway critical for auditory fear^22,24^. This pathway arises predominantly from the magnocellular division of MGB^25,26^ where its auditory magnocellular neurons, consistently described across species, show particular sensitivity to rapid temporal changes such as high rates of amplitude modulation (highAM)^27–32^. Similar to visual magnocellular channels, which also respond to rapid visual changes as major contributors to motion processing^29,33–35^, these auditory neurons may enable fast temporal coding of acoustic features relevant for threat processing. Such rapid temporal features have also been proposed as salient acoustic properties of alarm signals with evolutionary significance^36^. In humans, the magnocellular divisions of the auditory and visual thalamus have been histologically characterized and functionally linked to temporal processing in both sensory modalities^29,37–39^. Together, this raises the possibility that sounds with highAM could represent an auditory analogue of visual magnocellular-associated signals, engaging subcortical pathways for rapid emotional processing.

Building on this animal evidence, and on human findings in the visual domain, here we tested whether an auditory “low road” responsive to magnocellular-associated cues supports rapid threat processing in humans, paralleling the visual subcortical route. We combined behavioral, physiological, and neuroimaging measures across two complementary studies. During fear conditioning, participants detected human vocalizations that were modulated at either highAM or lowAM rates. These vocalizations acquired threatening value or remained neutral through Pavlovian association. Electroencephalography (EEG) and pupillometry were used to capture early neural and autonomic responses. Based on evidence in the visual domain^13–15^, we hypothesized that threat-related EEG and pupillary responses would emerge earlier for highAM, relative to lowAM voices^29,30,40^. In a separate sample of participants, we combined functional magnetic resonance imaging (fMRI) with diffusion-weighted MRI tractography to anatomically identify an auditory thalamo-amygdala pathway, and to assess its functional association with hemodynamic responses to highAM versus lowAM threat voices. If a subcortical auditory pathway to the amygdala operates through magnocellular-associated mechanisms specifically sensitive to highAM signals, we predicted that neural sensitivity to highAM threat cues would covary with individual variability in the fiber density of MGB-BLA connections. This approach allowed us to examine the functional and structural features of a subcortical auditory route for threat. Our findings support that the human brain may retain evolutionarily conserved subcortical pathways for threat processing across the sensory systems.

## Results

### Study 1. Identifying fast threat responses to highAM cues

#### Behavioral responses to threat-associated voices supported an effective fear conditioning

In a fear conditioning task, participants behaviorally indicated the location of male and female nonverbal voices, both modulated at highAM or lowAM rates (see Materials and Methods). During conditioning, one voice (CS+; a female voice for 50% of participants) was paired with an aversive white noise (US), while the other voice type remained unpaired (CS–; a male voice for the same 50% of participants). Before conditioning, the same voices were presented but with no aversive associations, enabling comparisons between physically identical stimuli that differed only in acquired emotional meaning (see Fig.1). Participants learned the association between the CS+ and the occurrence of the US, as revealed by CS-US contingency scores (main effect of CS: β = 1.960, SE = 0.12, *t*^(81)^ = 15.840, *p* < .001, r^2^ = .64), indicating that they predicted the occurrence of the US more strongly with CS+ stimuli compared to CS- stimuli. There was neither a significant main effect of AM (*p* = .182, r^2^ = .01), nor an AM x CS interaction (*p* = .290, r^2^ = .01). Participants judged emotional (CS+) stimuli as more negative compared to neutral (CS-), as revealed by valence ratings to voices analyzed on the subtracted values of the Conditioning minus the Pre-Conditioning phase (main effect of CS: β = 0.964, SE = 0.33, *t*^(108)^ = 2.890, *p* < .001, *r*^2^ = .07). There was neither a significant main effect of AM (*p* = .111, *r*^2^ = .02), nor a significant AM x CS interaction (*p* = .085, *r*^2^ = .03). Similarly, participants judged emotional (CS+) stimuli as more alerting than neutral (CS-) stimuli (main effect of CS on subtracted Conditioning minus Pre-Conditioning arousal scores: β = 0.893, SE = 0.24, *t*^(108)^ = 3.641, *p* < .001, *r*^2^ = .10). However, no significant main effect of AM (*p* = .771, *r*^2^ < .01) or AM × CS interaction was found (*p* = .138, *r*^2^ = .02).

**Figure 1.**
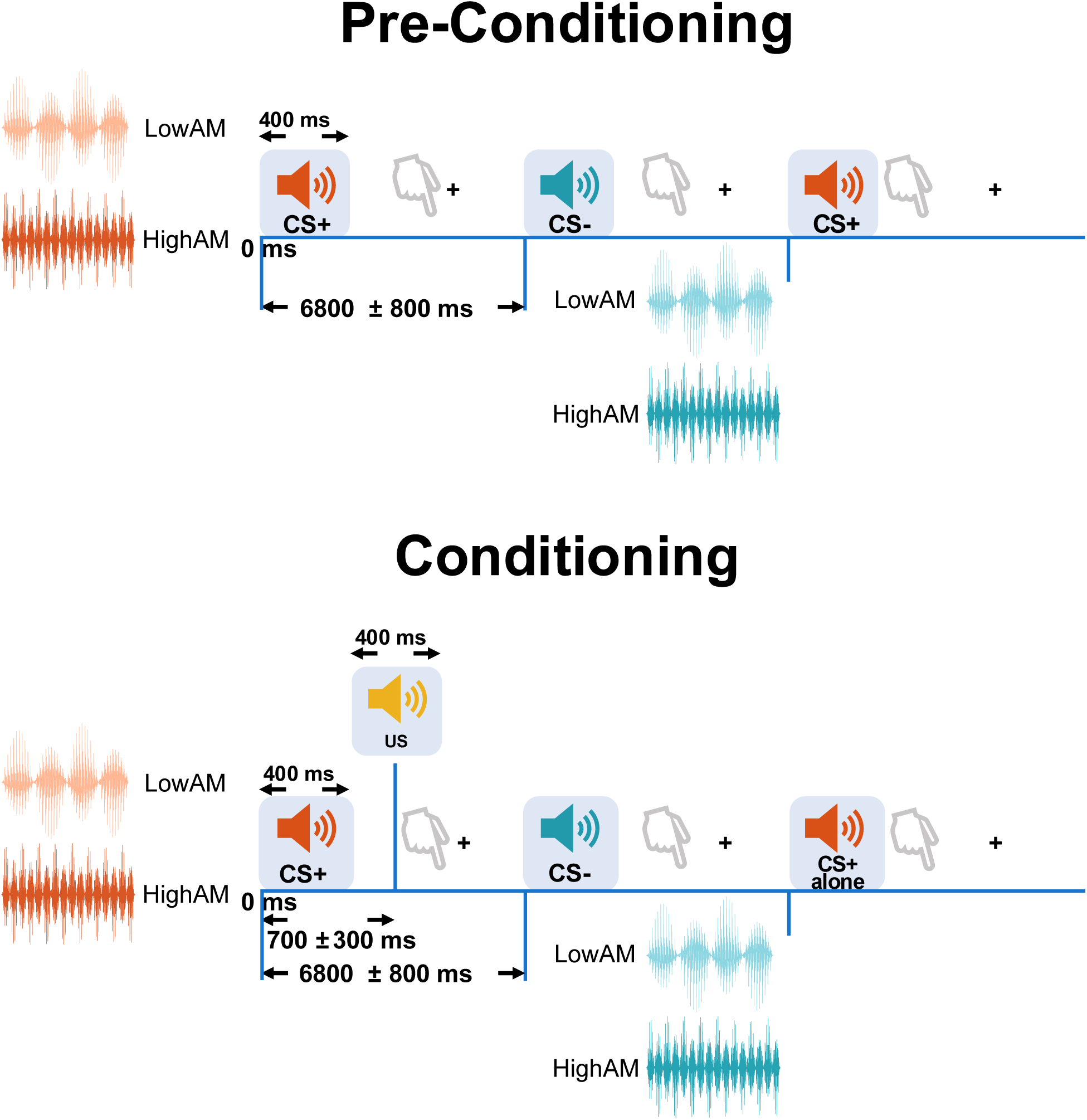
Experimental task. During conditioning, CS+ voices were paired with an unpleasant white noise (US), acquiring emotional value. In turn, CS- voices were never followed by the US, remaining neutral. Half of the CS+ trials were presented without the US (CS+ alone), allowing for posterior analysis of the physiological response in the absence of the US. During pre-conditioning, no voices were paired with the US, serving as control stimuli. Participants were asked to indicate whether each voice was presented on the left or right side, while maintaining central fixation. For half of the participants, the CS+ was a female voice, whereas the CS- was a male voice (and the opposite for the remaining sample). All voices were presented at either low or high amplitude modulation rates (lowAM, highAM, respectively).

Our linear mixed models on the subtracted response times (RTs) values for the Conditioning minus the Pre-Conditioning phase revealed that participants’ responses were slower for the CS+ compared to the CS- stimuli (main effect of CS: β = 0.067, SE = 0.02, *t*^(81)^ = 3.173, *p* < .01, *r*^2^ = .02) with no significant main effect of AM (*p* = .866, *r*^2^ = 5.45 × 10^-5^) or AM × CS interaction (*p* = .568, *r*^2^ = .001). A delayed response time of threat cues is consistent with increased defensive freezing-like processes commonly observed in fear conditioning^16,41,42^. Similar analyses with hit rate (HR) showed that participants were better at detecting the CS+ stimuli relative to the CS- (main effect of CS: β = 0.905, SE = 0.34, *t*^(54)^ = 2.648, *p* = .011, *r*^2^ = .03). An AM x CS interaction was also observed (β = -1.015, SE = 0.48, *t*^(54)^ = -2.100, *p* = .040, *r*^2^ = .02), suggesting that facilitated CS+ detection was stronger for highAM aversive sounds compared to lowAM aversive sounds. No significant main effect of AM (*p* = .267, *r*^2^ = .01) was observed. Behavioral plots are shown in Supplementary Fig. 1.

#### ERP and pupillary responses were increased for all threat-related voices at late time-windows

Our cluster-based permutation test on the ERPs in response to CS+ versus CS- voices (Conditioning minus Pre-Conditioning phase) revealed a late negative deflection that was larger for CS+ compared to CS-. This effect was maximal over fronto-central electrodes and spread toward parietal areas, and similar across highAM and lowAM sounds (highAM: 708 to 948 ms post-stimulus, *t*sum = 12,954.88, *p* = .007; lowAM: 622 to 996 ms post-stimulus, *t*sum = -17,956.20, *p* = .007; see Fig. 2A & Supplementary Fig. 2). Additionally, brain responses to highAM sounds were generally less negative than lowAM sounds within this late negative deflection over the same electrode regions (main effect of AM: 794 to 998 ms post-stimulus, *t*sum = 8,272.58, *p* = .010). No CS x AM interactions were found within this time window.

**Figure 2.**
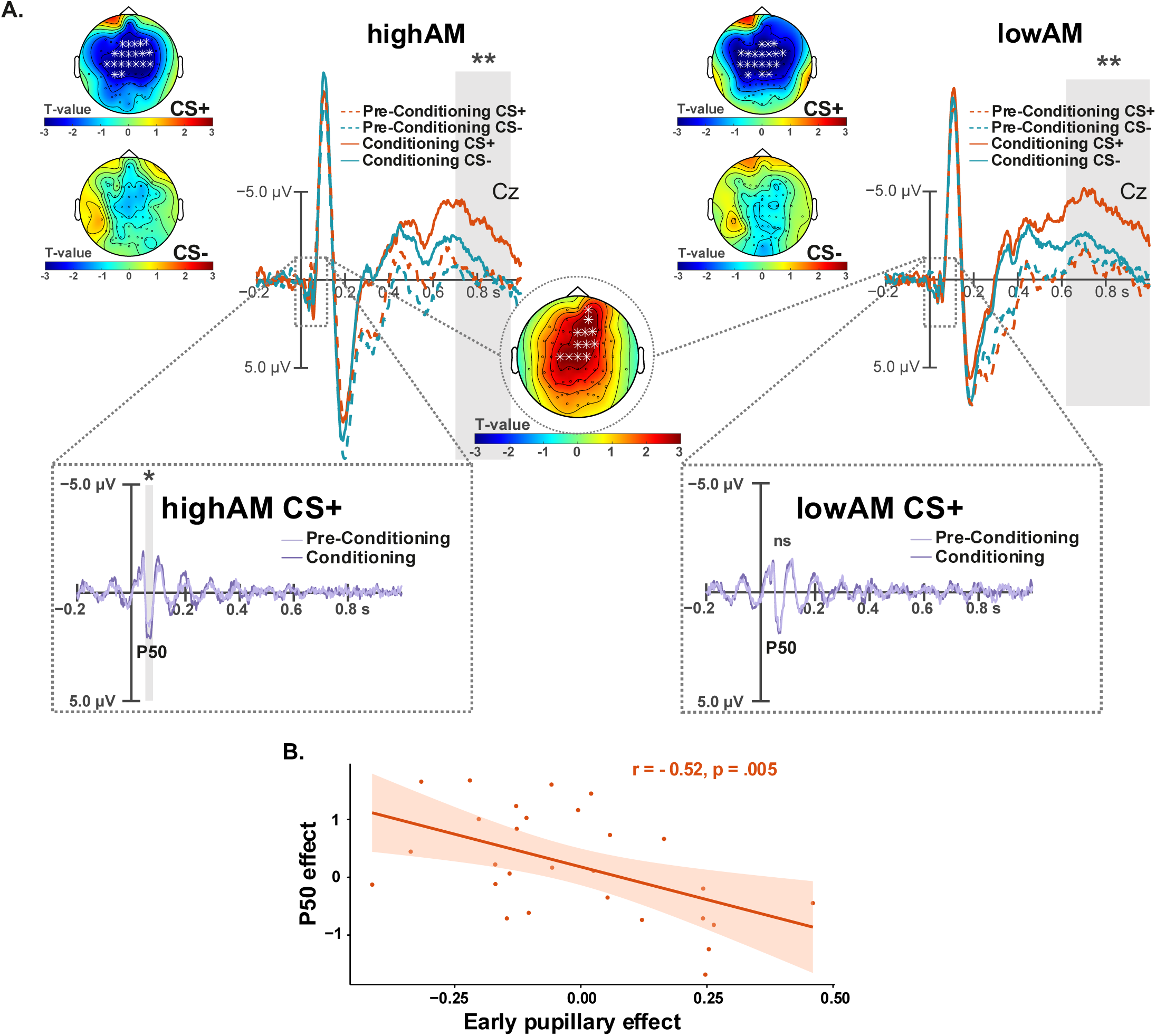
Neural responses to emotional (CS+) versus neutral (CS-) voices. (A) ERPs for highAM and lowAM voices (N=28). Top: Long-latency responses during Pre-Conditioning and Conditioning for both CS+ and CS- voices with topography maps for the significant (708-948 ms) window. Bottom: Middle-latency responses (P50) during Pre-Conditioning and Conditioning, and corresponding topography. (B) Association between early pupillary responses (<180 ms) and P50 amplitude (N=28). Shaded areas denote 95% confidence intervals. Grey areas indicate significant effects (* p ≤ .05, ** p ≤ .01). Abbreviations: RT = response time, ns = non-significant.

Finally, pupil diameter on the subtracted values (again with subtracted Conditioning minus Pre-Conditioning values) showed a late sustained increase in response to CS+ stimuli, relative to CS-, similarly for both highAM and lowAM stimuli (highAM: 740 to 1,500 ms post-stimulus, *t*sum = 246.04, *p* = .01; lowAM: 790 to 1,500 ms post-stimulus, *t*sum = 251.27, *p* = .01; Fig. 3, Supplementary Fig. 3A), with no statistical difference in this late window among AM conditions (Phase x CS x AM: *t*sum = -4.96, *p* = 1). A late and sustained increase in ERP and pupillary responses supports an effective attention and arousal response associated with conditioning in participants.

**Figure 3.**
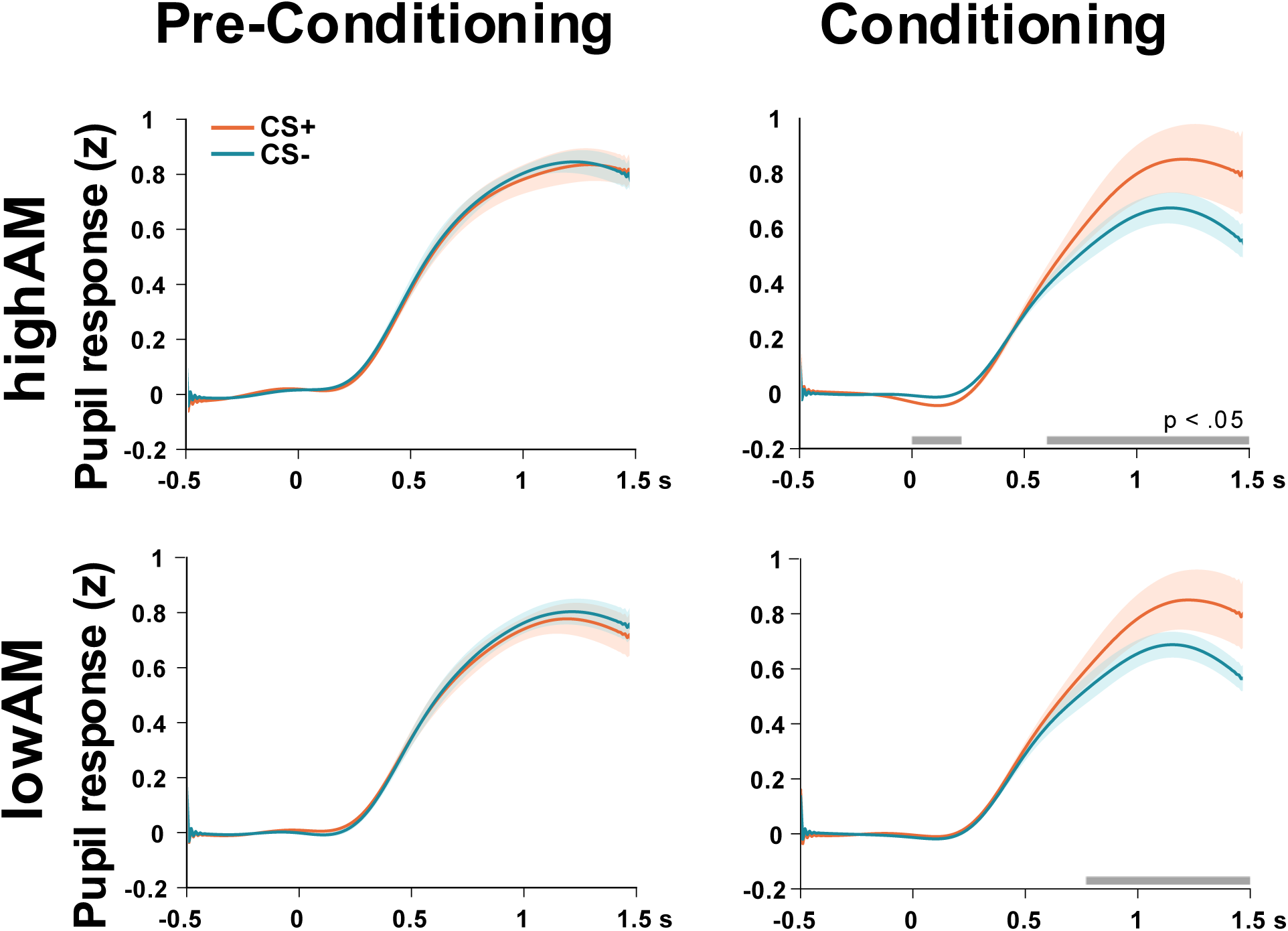
Pupillary responses to the sounds. (Top row) Pupil responses to emotional (CS+) and neutral (CS-) voices for each condition and conditioning phase. Shaded areas represent the standard error of the mean in all figures.

#### Early threat responses were selectively associated with highAM voices

When evaluating conditioning on the long-latency ERPs (Fig. 2A), we identified an early Phase x CS x AM significant cluster spanning from 30 to 78 ms over fronto-central electrodes (cluster-level *p* = .002, *t*sum = 20,331.10; 10,000 permutations), suggesting that conditioned CS+ sounds elicited an early increase in the auditory response, particularly when presented at highAM, relative to lowAM. For a better approximation of this early effect (as cluster-based statistics may overestimate the timing), we further explored the auditory ERP by applying standardized filters to the EEG signal, so as to isolate the middle-latency responses (Rentzsch et al.^43^; Fig. 2A). The analysis, again implemented directly on the difference wave for Conditioning minus Pre-Conditioning, confirmed a clear amplitude increase of the P50 component that was only present for highAM voices (AM effect: β = -0.230, SE = 0.11, t^(82)^ = -2.187, p = .031, r^2^ = .03). This increased response to highAM versus lowAM voices occurred irrespective of CS type (CS effect: p = .99) but emerged selectively during the threat-conditioning phase (Conditioning versus Pre-Conditioning), consistent with a phase-specific threat response.

Remarkably, our cluster-based analysis of the pupillary response also revealed an early reduction in pupil diameter that was specific to the Conditioning phase (versus Pre-Conditioning) and restricted to highAM CS+ voices. This early effect spanned the first 180 ms post-stimulus (CS x AM: 0 to 180 ms post-stimulus, p = .024, tsum = -45.75; Fig. 3 & Supplementary Fig. 3B).

#### Early neural and pupillary responses to highAM threat voices were mutually associated

To further examine the relationship between early neural and autonomic responses to highAM aversive voices, we computed Pearson correlations between individual pupil diameter and P50 amplitude values, extracted from the 0 - 220 ms time-window where main effects were observed (each value corresponding to a three-way difference: Conditioning minus Pre-Conditioning, CS+ minus CS- and high AM minus low AM). Pupil responses showed a negative correlation with auditory P50 amplitude (*r* = -.52, *p* = .005, 95% *CI*: -0.75, -0.18), indicating that stronger auditory responses to highAM threat voices were associated with greater early pupil constriction (Fig. 2B). This suggests a functional association between auditory and pupillary responses to highAM aversive stimuli, potentially reflecting coordinated activity during the initial stages of threat processing. Complementary analyses examining the relationship between early ERP responses and detection performance are reported in the SI Results.

### Study 2. Associating an MGB-BLA pathway with threat-related responses to highAM cues

#### Amygdala response to highAM versus lowAM threat voices covaried with fiber density of direct MGB-BLA connections

Combining fMRI and diffusion-weighted tractography, we examined whether interindividual variability in the structural strength of direct auditory thalamo-amygdala connections predicted threat-related response to highAM versus lowAM threat cues in the amygdala (see SI Materials and Methods). No association was observed between fiber density of the left MGB-BLA tract and brain responses to threat. In turn, fiber density of the right MGB-BLA tract was not associated with brain activity for the main AM effect of interest (highAM versus lowAM). However, fiber density of this right tract covaried with hemodynamic activity in right amygdala for the AM x CS interaction ([highAM CS+ > highAM CS-] – [lowAM CS+ < lowAM CS-]) (MNI coordinates: 28, -6, -22; kE = 80; T = 4.59; pFWE-SVC < .001). In contrast, no such association was observed for the reverse interaction ([highAM CS+ < highAM CS-] – [lowAM CS+ > lowAM CS-]). Post-hoc contrasts for highAM emotional cues [highAM CS+ > highAM CS−] confirmed a positive relationship between fiber density of the tract and right amygdala response (MNI coordinates: 26, -6, -20; kE = 87; T = 4.23; pFWE-SVC = .002; Fig. 4 and Supplementary Table 1). This association indicates that individuals with stronger anatomical connectivity between the right MGB and amygdala exhibited greater ipsilateral amygdala response to highAM threat voices. Importantly, this effect was selective to highAM cues, as no comparable association was observed for the lowAM emotional contrast [lowAM CS+ > lowAM CS−] (see Fig. 4 and Supplementary Table 1). For completeness, a whole-brain analysis (FWE-corrected) was also computed for all contrasts of interest, with no significant anatomo-functional associations. Together, these results provide convergent evidence that a direct auditory thalamo-amygdala pathway is preferentially tuned to highAM threat cues, consistent with a fast “low road” for threat associated with magnocellular processing.

**Figure 4.**
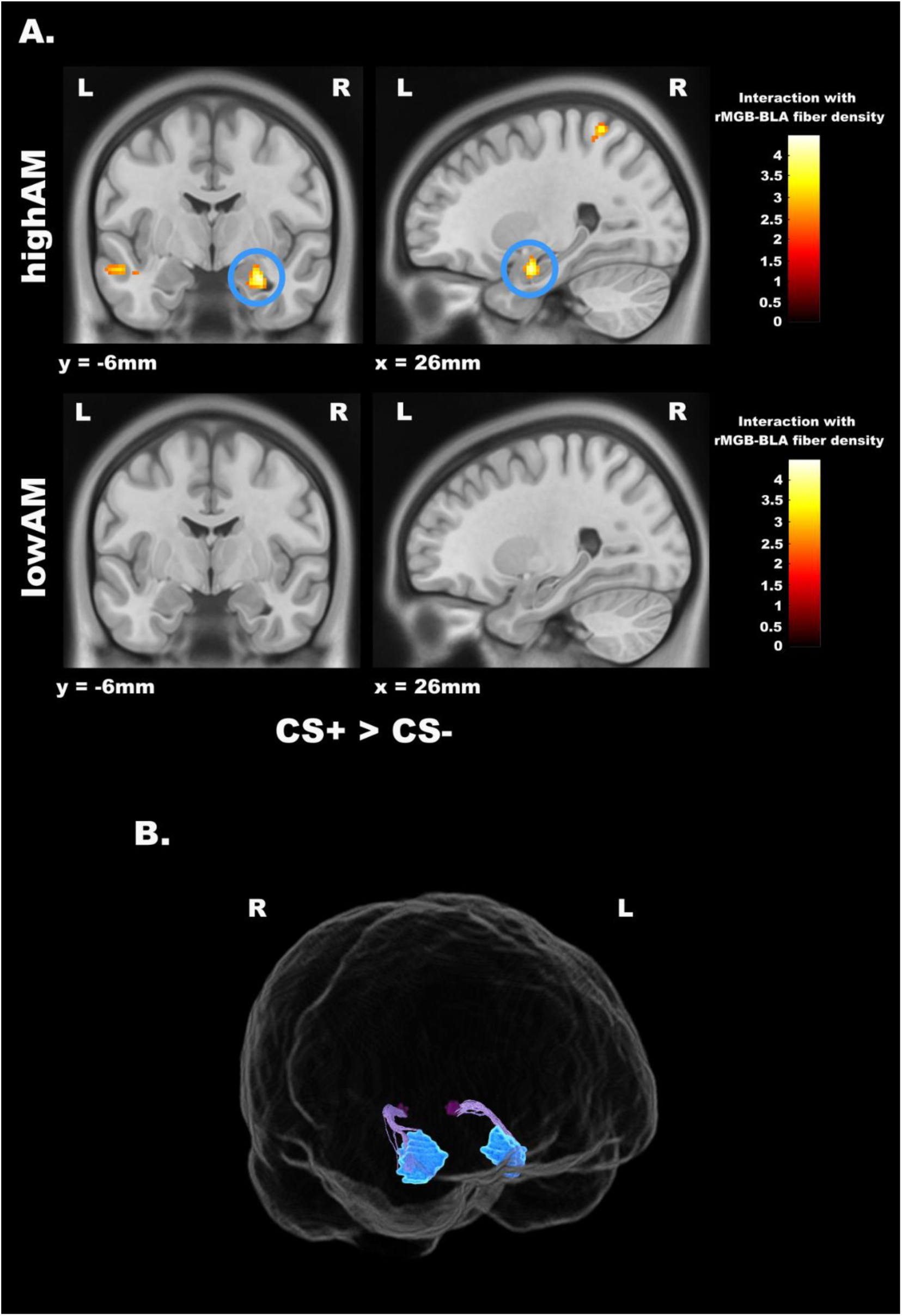
Association between hemodynamic responses to highAM threat and anatomical connectivity of an auditory thalamo-amygdala pathway. (A) Functional MRI combined with tractography analysis of connections between the auditory thalamus (MGB) and the basolateral amygdala (BLA). Top: Right amygdala activity for highAM CS+ > highAM CS- voices that covaried with fiber density of the right MGB-BLA pathway. Bottom: No association for lowAM CS+ > lowAM CS-. All contrasts are displayed at *p* < .01 for visualization. (B) 3D probabilistic tractography reconstruction of the MGB-BLA pathway in a representative subject. Fibers are seeded from the MGB (dark purple) and terminate in the BLA (blue), with streamlines shown in light purple. Abbreviations: L = left, R = right.

In sum, threat conditioning modulated behavior, neural, and autonomic responses^44–47^, but only highAM voices triggered rapid neural and pupillary effects, consistent with fast inputs to the amygdala. Remarkably, the structural strength of a direct right MGB-BLA pathway predicted right amygdala sensitivity to highAM (versus lowAM) threatening voices. These results are compatible with a fast subcortical auditory pathway for threat in humans that may be preferentially engaged by magnocellular-associated acoustic signals.

## Discussion

Our findings provide convergent physiological and neuroimaging evidence for a fast and direct auditory route to the amygdala that is preferentially sensitive to rapidly modulated (highAM) threat cues. This pathway may thus represent an auditory analogue of the visual magnocellular “low roads” proposed to mediate emotional processing in humans^2,9,48^. Through Pavlovian conditioning, we manipulated both the aversive meaning and the acoustic AM rate of voices, allowing us to isolate physiological responses linked to threat processing for highAM versus lowAM sounds. By extending animal evidence to humans, our results provide a framework for understanding evolutionarily conserved subcortical mechanisms of emotion processing across modalities.

Using EEG and pupillometry, we identified early neural and pupillary responses to threat, and found that these responses depended on highAM, rather than lowAM rates, whereas later responses were comparable across all acoustic conditions. Given that highAM sounds have been associated with magnocellular-related inputs in the auditory system^29–31,38^, our results provide empirical support for rapid emotion signals that may rely on magnocellular-associated acoustic cues^8,11–13^. This is comparable to fast visual magnocellular-related signals preferentially conveyed by low spatial-frequency stimuli during emotion processing^11,13,14^. Indeed, magnocellular auditory neurons are considered homologous to magnocellular neurons in the visual system, as both respond to similar parameters, including low spatial frequencies as well as rapid amplitude and frequency changes over time^29,30,32,37^. These results may thus be consistent with rapid transmission of affective signals through direct auditory thalamo-amygdala projections previously described in rodents and non-human primates^49–51^, which may be sensitive to magnocellular-associated inputs, similarly as in vision.

Despite differences in visual and auditory latencies across sensory modalities, our observed increase in amplitude of the auditory P50 component (48-78 ms post-stimulus) for highAM versus lowAM voices is compatible with early human amygdala responses previously described for visual low spatial-frequency (LSF) fearful faces^14^, where rapid amygdala input may transiently enhance cortical excitability for all incoming stimuli before later cortical discrimination^8,52,53^. This is also consistent with amygdala responses preceding those in visual cortex for fearful faces under similar LSF conditions^13^. More directly, our findings converge with ultrafast responses in the rodent auditory cortex when threatening sounds are processed through the subcortical route to amygdala^54^. We note that this P50 effect occurred similarly for CS+ and CS− stimuli, but was specific to the conditioning phase, and therefore associated with threat, possibly reflecting generalized arousal mediated by the amygdala.

In parallel with the neural effects, our observed early pupillary constriction within the first 180 ms was also specific to highAM aversive voices. Although further research is needed, these findings may also suggest a contribution of magnocellular-associated inputs on early autonomic responses, presumably through amygdala circuitry. The basolateral amygdala projects to the central amygdala nucleus, which can in turn influence sympathetic arousal-related responses to threat^4^ through projections to locus coeruleus (LC), mediating pupil dilation, but also parasympathetic responses via the periaqueductal gray to the Edinger-Westphal (EW) nucleus, mediating pupil constriction^1,55–57^. Differential engagement of distinct circuitries could underlie the rapid pupil dynamics observed for highAM threat, possibly reflecting transient subcortical activity.

Remarkably, a temporal overlap and linear association between enhanced P50 amplitude and early pupil constriction suggested a potential functional link between neural and autonomic responses during the initial stages of threat processing. Thus, participants with stronger P50 amplitude increases to highAM aversive voices also showed greater pupil constriction, suggesting that both responses may originate from coordinated rapid mechanisms supporting adaptive fear behavior. Together, early responses selective to highAM emotional sounds are compatible with rapid threat processing, potentially mediated by fast temporal cues associated with magnocellular processing^1,21,54^. In line with this interpretation, highAM threat cues were also associated with enhanced detection performance, aligning with facilitated detection of threat under these acoustic conditions.

Finally, our multimodal imaging approach allowed us to identify a direct pathway connecting auditory thalamus (MGB) with amygdala (BLA) and showed that interindividual variability in the structural strength of this tract covaried with functional amygdala responses to highAM versus lowAM threat voices. This supports a preferential sensitivity of the pathway for fast temporal threat cues that are associated with magnocellular processing, consistent with our physiological findings. Moreover, this anatomo-functional association was right-lateralized, aligning with prior visual work implicating the right amygdala in rapid or unconscious threat processing particularly for low-spatial-frequency stimuli^17,18,58^. These findings converge with animal work demonstrating that direct MGB-BLA projections are involved in auditory fear processing and learning^20,22,25,26,50^, and support their preferential sensitivity to magnocellular-associated acoustic signals^25,27,50,59^.

Although previous work in humans has sought to identify an auditory “low road” using functional connectivity approaches^60,61^, this has yet to provide explicit anatomical evidence for MGB-amygdala association with auditory fear. Here, we identify a structurally defined human pathway and characterize its functional association with fast temporal threat cues, advancing current understanding of human subcortical emotion processing.

In human speech, lowAM cues typically convey fine-grained linguistic information, such as syllabic or prosodic cues essential for semantic encoding^62^. Accordingly, processing of lowAM threat cues may preferentially recruit slower, cortically mediated inputs to the amygdala, analogous to parvocellular channels in vision^8,11,29,36,63^. Conversely, highAM cues may preferentially recruit coarse acoustic processing and correlate with sound “roughness”, a hallmark of screams and alarm signals that strongly engage the amygdala^36,64^. Thus, a rapid prioritization of highAM threat cues likely reflects their biological salience as warning signals, enabling fast sensory-autonomic coupling^8,10,36^ presumably through evolutionarily conserved thalamo-amygdala circuits. Such circuits may be deeply rooted in mammalian evolution, with homologous MGB-amygdala projections supporting rapid defense responses across species^22,26^.

Importantly, an association of the MGB-BLA tract with highAM cues suggests preferential sensitivity of this tract for amplitude modulations around 40 Hz. However, this does not necessarily imply that thalamo-amygdala connections are never recruited by lower-rate amplitude-modulated cues, or other acoustic parameters or conditions. Indeed, MGB-BLA circuitry in non-human species is known to contribute to numerous threat-related functions^20,25,54^. Similarly, the visual “low road” in humans may be tuned to distinct stimulus features beyond lower or higher spatial frequencies^65^, including subliminal threat signals^9,14,16^, peripherally presented stimulation^66,67^, or even auditory emotional cues^68,69^. Future work should disentangle the contribution and specificity of different subcortical routes across affective and sensory domains, clarifying how evolution has shaped threat processing in the human brain.

Finally, the early emotion effects observed for highAM voices may not be explained by physical differences between highAM and lowAM stimuli. Although the faster temporal envelope of highAM sounds could in principle produce earlier peripheral or cortical auditory transduction (see Fig. 1), this explanation is unlikely, as no ERP differences emerged between highAM and lowAM voices during pre-conditioning, before emotional associations took place. This rules out trivial acoustic confounds and supports early threat-related modulation (see SI Materials and Methods and SI Results). Moreover, because EEG does not allow localization of amygdala activity, we cannot fully exclude the contributions from cortical^50,70^, purely thalamic, or brainstem generators^1,71,72^, without amygdala contribution in our physiological effects. However, our imaging results support thalamo-amygdala contributions to threat processing under the same acoustic conditions, and align with extensive animal evidence showing a critical role of MGB-BLA projections in affective function^20,25,54^.

In sum, our findings provide convergent empirical evidence for an auditory “low road” in humans that is responsive to magnocellular-associated features, potentially mirroring the subcortical routes long described in vision. This circuitry may reflect an evolutionarily conserved mechanism enabling rapid processing of coarse acoustic threat cues^36,64,73^. Visual and auditory magnocellular systems share response properties and likely derive from common ontogenetic origins^28,29,74^, supporting the idea of ancestral common mechanisms for fast affective processing across modalities. Moreover, given that dysfunction of the visual route and associated processing have been linked to psychiatric conditions such as anxiety, schizophrenia, or autism^2,75^, our findings may therefore contribute to understanding early sensory-affective disturbances in the auditory domain^76^. By revealing a subcortical auditory route sensitive to fast temporal inputs, our work may also bridge animal and human evidence, while contributing to a unified cross-modal framework for emotion processing.

## Materials and Methods

### Study 1: EEG and pupillometry

We conducted this study to test whether highAM voices elicit fast neural and autonomic signatures of fear, relative to lowAM voices. This would be consistent with a fast route for auditory threat processing in humans sensitive to magnocellular-associated features.

### Subjects

Thirty-two healthy volunteers were mostly recruited among University of Barcelona students. All were right-handed, with normal hearing and no previous neurological or psychiatric disorders. Four participants were excluded from the analysis due to low signal-to-noise ratio on the electrophysiological data. Thus, the final sample consisted of twenty-eight participants (12 females; age range 18-31, mean age = 22 years). All procedures were approved by the ethics committee of the University of Barcelona, in accordance with the Declaration of Helsinki (2024). All volunteers provided written informed consent and received monetary compensation for their participation.

### Stimuli

Voices are phylogenetically relevant stimuli and may serve as biologically significant signals^77–79^ to which the amygdala is especially sensitive^19^. Two 400-ms excerpts of female and male non-verbal vocal utterances with neutral prosody, extracted from the Montreal Affective Voices database^80^, served as conditioned stimuli (CS). They were digitally modulated in amplitude at high (40 Hz) and low (10 Hz) amplitude modulation (AM) rates using MATLAB (MathWorks, R2019b) with a 100% AM depth and 10 ms rise and fall ramps, resulting in 4 different stimuli. Voices and AM rates were selected based on previous literature^36^ and preliminary ratings from 6 independent participants, and those rated as emotionally neutral were chosen. HighAM and lowAM stimuli were designed to differentially engage auditory fast (magnocellular-like) and slow pathways, respectively^29^. Stimuli were delivered at 80 ± 1 dB SPL on average across participants, while ensuring that they were clearly audible to them. They were presented binaurally and lateralized towards the right or left by manipulating interaural time differences (ITD; Arnal et al.^36^), Additionally, a 400-ms binaural burst of an unpleasant, but not painful, loud white noise served as unconditioned stimulus (US) individually calibrated (mean = 103 ± 7 dB SPL). All stimuli were delivered through headphones (Sennheiser HD 558) using Psychtoolbox (Version 3.0.14).

### Procedure and task

The experiment was conducted in a dimly lit room, with participants seated in front of a screen, and their heads stabilized in a chinrest positioned at a distance of 60 cm. Participants were instructed to respond, as quickly and accurately as possible, on which side they heard the voice (left or right, by pressing a keyboard button with their right index or middle finger, respectively), while maintaining their gaze on a central fixation cross. Voices were presented in trials of 6800 ± 800 ms duration (Domínguez-Borràs et al.^41^; see Fig. 1). The task followed a classical fear-conditioning paradigm, which depends on sensory inputs to the amygdala^6^. Conditioning allowed us to confer aversive meaning to acoustic stimuli while independently manipulating their acoustic parameters. Thus, we could compare brain and pupil responses to identical stimuli across conditioning phases (after and before conditioning), as well as across stimulus categories differing in their acquired affective meaning (conditioned/emotional or non-conditioned/neutral), ensuring that emotional effects were dissociated from physical stimulus properties.

The task consisted of a Pre-Conditioning, a Conditioning (Fig. 1), and a short Extinction phase (not analyzed) to remove the emotional value of the stimuli at the end of the experiment. During Conditioning, participants were presented with both highAM and lowAM voices, which were either paired (CS+) or unpaired (CS-) with the unpleasant loud white noise (US). Stimulus assignment as either CS+ or CS- was based on voice gender and counterbalanced across participants (i.e., 50% were presented with highAM and lowAM CS+ male voices, and with highAM and lowAM CS- female voices, and vice versa for the other 50%). The CS-US pairing conferred aversive emotional (fear-related) significance to the CS+, whereas the CS- remained neutral.

During Pre-Conditioning, the same CS stimuli were presented without the US, allowing comparisons among identical stimuli before and after they acquired emotional value. To isolate physiological CS+ responses uncontaminated by the US, conditioning was conducted with a 50% partial reinforcement schedule, such that only half of the CS+ trials were presented with the US, resulting in 50% of CS+ stimuli with no US (CS+ alone; see Fig.1)^47,81^. The final Extinction phase was identical to Pre-Conditioning, but its sole purpose was to allow the stimuli to lose affective value. Thus, there were four conditions of interest (analyzed) in the Conditioning phase (*CS+ alone highAM*, *CS+ alone lowAM*, *CS- highAM*, *CS- lowAM*), and four control conditions in the Pre-Conditioning phase (*CS+ highAM*, *CS+ lowAM*, *CS- highAM*, *CS- lowAM*). The number of trials and their distribution across conditions are detailed in SI Materials and Methods.

Voices were presented in pseudo-random order, ensuring no more than two consecutive repetitions of the same stimulus to prevent habituation^45^. Voices were presented ensuring that an equal number of voices was perceived on each side. Trials without participants’ responses within 2 s after voice onset were counted as misses.

At the end of each block of the Conditioning phase, participants rated on a scale from 1 (never) to 5 (always), how likely each voice of the experiment was to be followed by the US, to assess their awareness of the CS-US contingency. Finally, at the beginning and end of the Conditioning phase, participants rated all voices for valence and arousal on scales from 1 (very positive/low arousal) to 5 (very negative/high arousal).

### Electrophysiology

EEG was recorded at 500 Hz from 64 scalp electrodes (10–20 system; Neuroscan SynAmps RT). Horizontal and vertical EOGs were monitored, and impedances were kept below 10 kΩ. Data were bandpass-filtered (0.1–100 Hz for long-latency and 10–200 Hz for middle-latency responses), re-referenced to the nose, and corrected for ocular artifacts using independent component analysis (SOBI). Epochs (−200–1000 ms) were baseline-corrected and trials exceeding ±75 μV were rejected. Details on filtering and preprocessing are provided in SI Materials and Methods.

A mass-univariate nonparametric cluster-based permutation analysis was conducted^82,83^ with Fieldtrip^84^ on ERP amplitudes using a two-dimensional (time x electrode) approach (individual ERP data to compute main effects; individual ERP difference data across conditions to compute the interaction). Clusters were defined via Delaunay triangulation requiring at least two neighboring electrodes. One-tailed dependent t-tests were applied to the amplitudes at each time x electrode point. P-values were estimated by summing t-values of adjacent significant samples (*p* < .05; two-tailed), using 10,000 Monte Carlo randomizations. Corrected p-values below .025 (one-tailed) were considered significant. For each significant cluster, we report its temporal spread, cluster statistic, and p-value.

Effects of conditioning, as a measure of the emotional response, were assessed with a CS (CS+ versus CS-) x Phase (Conditioning versus Pre-Conditioning) interaction on difference waves of pairs of conditions. This subtraction-method approach reduced complexity of the statistical models and controlled for physical differences between the CS voices. Thus, we subtracted the CS+ condition in the Conditioning phase versus the same CS+ condition in the Pre-Conditioning phase, and compared this difference to the equivalent subtraction for CS-. Similarly, to test how amplitude modulation (highAM versus lowAM) impacted the emotional response, we computed a Phase x CS x AM interaction with an analogous subtraction approach ([Conditioning CS+ alone highAM minus Pre-Conditioning CS+ highAM] minus [Conditioning CS- highAM minus Pre-Conditioning CS-highAM], and then statistically comparing this double subtraction with the equivalent computation for lowAM). To further explore the early responses observed with this data-driven analysis, we analyzed the middle-latency P50 response (40-80 ms post-stimulus) at electrode Cz^85^. Phase x CS x AM effects were further assessed with the same subtraction approach using a linear mixed model (LMM) including CS, AM and their interaction as fixed effects, and Participant as random effect, using the lmer function^86^ of RStudio (R Core Team, 2024). LMMs were chosen for robustness to moderate violations of normality in some of our measures^87^. All analyses were done after confirming all the model assumptions were met.

### Pupillometry

Pupil diameter was recorded from the left eye using an EyeLink 1000 Desktop Mount (SR Research; sampling rate = 1000Hz) while participants maintained their gaze on a central black fixation cross. The eye tracker was calibrated at the beginning of the experiment and during breaks. Preprocessing and analysis were conducted with custom written MATLAB scripts and Fieldtrip.

Missing data and blinks were detected and linearly interpolated^88^ were padded. Blinks less than 250 ms apart were merged into a single blink. Signal was bandpass filtered (0.05 to 4 Hz third-order Butterworth; Binda et al.^89^) with blinks and saccades removed via linear regression^88,90^. Residual pupil time series was z-scored and resampled to 100 Hz. Trials were epoched from −0.5 to +1.5 post-stimulus, baseline corrected (with a 500-ms pre-stimulus) and averaged per condition.

Nonparametric permutation statistics (10,000 Monte Carlo iterations) were employed as in the EEG analyses, except clusters here were defined along the temporal dimension. Main and interaction effects of CS, Phase, AM were assessed analogously, computing t-values between subtracted conditions and thresholding them at *p* < .05 (two-tailed dependent test). Finally, pupillary responses were correlated with early auditory ERP amplitudes using Pearson’s correlation (two-tailed). All the model assumptions were verified to be met.

### Behavior

Response times (RTs) for correct trials (errors and omissions were below 2%) were computed per subject and condition (Supplementary Fig. 1D). Hit rates (HR) were analyzed per condition to assess stimulus detection (Supplementary Fig. 1E). For valence and arousal ratings, we computed difference scores by subtracting Pre-Conditioning from Conditioning ratings. Contingency ratings were averaged across all blocks during Conditioning. Valence, arousal and contingency ratings were analyzed separately for each of the four voice conditions (see Supplementary Fig. 1A-C). After confirming all the model assumptions were met, LMMs were applied to behavioral measures using again CS, AM, and CS x AM) as fixed effects, with Participant as a random effect.

### Study 2: functional MRI and diffusion tractography

With this study, we tested whether amygdala responses to highAM threat cues covaried selectively with the strength of MGB-BLA anatomical connectivity, supporting a subcortical route for auditory threat associated with magnocellular-related cues. An independent sample of thirty-one healthy volunteers (18 females; age range 18–39, mean = 24.4 years) participated in an fMRI and diffusion-weighted MRI study using an adapted version of the conditioning task from Study 1 (see SI Materials and Methods). Procedures were identical except that the US consisted of an aversive white noise combined with some aversive images taken from a standardized database to reinforce the aversive effect of the US in the scanner (see SI Materials and Methods). In addition, stimulus assignment as either CS+ or CS- was based on the source location of each voice instead of the gender of the stimulus voice (i.e., 50% were presented with highAM and lowAM CS+ left-sided voices, and with highAM and lowAM CS- right-sided voices). MRI data were acquired on a 3T Philips Ingenia CX scanner with a 32-channel head coil, including high-resolution T1-weighted structural scans, whole-brain fMRI, and multi-shell diffusion-weighted imaging. Functional data were preprocessed and analyzed in SPM12.

Diffusion data were preprocessed in MRtrix3 with standard denoising, motion and distortion correction, multi-shell multi-tissue constrained spherical deconvolution, bias correction, and normalization. Anatomical MGB and BLA ROIs were defined using FreeSurfer segmentations. Probabilistic tractography of the MGB-BLA pathway was conducted in MRtrix3 using ACT and SIFT2. Fiber bundle capacity (FBC) values were extracted as quantitative indices of MGB-BLA connectivity strength, reflecting streamline fiber density. For clarity, we refer to these values as fiber density throughout the text. Finally, to assess the relationship between interindividual MGB-BLA anatomical connectivity and functional responses to highAM versus lowAM threat cues, individual right and left fiber density values were entered as covariates in a flexible factorial modeled with the Conditioning - Pre-Conditioning contrasts for each condition of interest. The main contrasts of interest tested the condition x covariate interaction for the AM x CS effect ([highAM CS+ > highAM CS-] – [lowAM CS+ < lowAM CS-]), as well as for the post-hoc contrasts [HighAM CS+ > HighAM CS-] and [LowAM CS+ > LowAM CS-], (see SI Materials and Methods). For completeness a whole analysis was also computed for the contrasts of interest.

## Acknowledgments

This work was supported by the European Research Council (ERC) under the European Union’s Horizon 2020 research and innovation program (Grant agreement No.101088954), the Spanish Ministry of Science and Innovation project (PID2020-116311GA-I00), the María de Maeztu Center of Excellence (MDM-2017-07-29-20-2), as well as the Generalitat de Catalunya (SGR2021-00356).

## References

1. LeDoux, J. Emotional networks and motor control: A fearful view. Prog. Brain Res. 107, 437–446 (1996).

2. McFadyen, J., Dolan, R. J. & Garrido, M. I. The influence of subcortical shortcuts on disordered sensory and cognitive processing. Nat. Rev. Neurosci. 21, 264–276 (2020).

3. Carr, J. A. I’ll take the low road: the evolutionary underpinnings of visually triggered fear. Front. Neurosci. 9, 414 (2015).

4. Janak, P. H. & Tye, K. M. From circuits to behaviour in the amygdala. Nature 517, 284–292 (2015).

5. Ledoux, J. E. & Brown, R. A higher-order theory of emotional consciousness. Proc. Natl. Acad. Sci. U. S. A. 114, E2016–E2025 (2017).

6. LeDoux, J. E. Emotion circuits in the brain. Annu. Rev. Neurosci. 23, 155–184 (2000).

7. Phelps, E. A. & LeDoux, J. E. Contributions of the amygdala to emotion processing: From animal models to human behavior. Neuron 48, 175–187 (2005).

8. Vuilleumier, P. How brains beware: Neural mechanisms of emotional attention. Trends Cogn. Sci. 9, 585–594 (2005).

9. Tamietto, M. & De Gelder, B. Neural bases of the non-conscious perception of emotional signals. Nat. Rev. Neurosci. 11, 697–709 (2010).

10. Domínguez-Borràs, J. & Vuilleumier, P. Amygdala function in emotion, cognition, and behavior. Handb. Clin. Neurol. 187, 359–380 (2022).

11. Vuilleumier, P., Armony, J. L., Driver, J. & Dolan, R. J. Distinct spatial frequency sensitivities for processing faces and emotional expressions. Nat. Neurosci. 6, 624–631 (2003).

12. McFadyen, J. Investigating the Subcortical Route to the Amygdala Across Species and in Disordered Fear Responses. J. Exp. Neurosci. 13, 1179069519846445 (2019).

13. Méndez-Bértolo, C. et al. A fast pathway for fear in human amygdala. Nat. Neurosci. 19, 1041–1049 (2016).

14. Wang, Y. et al. Rapid Processing of Invisible Fearful Faces in the Human Amygdala. J. Neurosci. 43, 1405–1413 (2023).

15. Pourtois, G., Dan, E. S., Grandjean, D., Sander, D. & Vuilleumier, P. Enhanced extrastriate visual response to bandpass spatial frequency filtered fearful faces: Time course and topographic evoked-potentials mapping. Hum. Brain Mapp. 26, 65–79 (2005).

16. Celeghin, A., de Gelder, B. & Tamietto, M. From affective blindsight to emotional consciousness. Conscious. Cogn. 36, 414–425 (2015).

17. Pegna, A. J., Khateb, A., Lazeyras, F. & Seghier, M. L. Discriminating emotional faces without primary visual cortices involves the right amygdala. Nat. Neurosci. 8, 24–25 (2005).

18. Burra, N., Hervais-Adelman, A., Celeghin, A., de Gelder, B. & Pegna, A. J. Affective blindsight relies on low spatial frequencies. Neuropsychologia 128, 44–49 (2019).

19. Diano, M., Celeghin, A., Bagnis, A. & Tamietto, M. Amygdala Response to Emotional Stimuli without Awareness: Facts and Interpretations. Front. Psychol. 7, 2029 (2016).

20. Bordi, F. & LeDoux, J. E. Response properties of single units in areas of rat auditory thalamus that project to the amygdala - II. Cells receiving convergent auditory and somatosensory inputs and cells antidromically activated by amygdala stimulation. Exp. Brain Res. 98, 275–286 (1994).

21. Quirk, G. J., Repa, J. C. & LeDoux, J. E. Fear conditioning enhances short-latency auditory responses of lateral amygdala neurons: Parallel recordings in the freely behaving rat. Neuron 15, 1029–1039 (1995).

22. Shinonaga, Y., Takada, M. & Mizuno, N. Direct projections from the non-laminated divisions of the medial geniculate nucleus to the temporal polar cortex and amygdala in the cat. J. Comp. Neurol. 340, 405–426 (1994).

23. Aggleton, J. P., Burton, M. J. & Passingham, R. E. Cortical and subcortical afferents to the amygdala of the rhesus monkey (Macaca mulatta). Brain Res. 190, 347–368 (1980).

24. LeDoux, J. E., Sakaguchi, A. & Reis, D. J. Subcortical efferent projections of the medial geniculate nucleus mediate emotional responses conditioned to acoustic stimuli. J. Neurosci. 4, 683–98 (1984).

25. LeDoux, J. E., Farb, C. & Ruggiero, D. A. Topographic organization of neurons in the acoustic thalamus that project to the amygdala. J. Neurosci. 10, 1043–1054 (1990).

26. Romanski, L. M. & LeDoux, J. E. Equipotentiality of thalamo-amygdala and thalamo-cortico-amygdala circuits in auditory fear conditioning. J. Neurosci. 12, 4501–4509 (1992).

27. De La Mothe, L. A., Blumell, S., Kajikawa, Y. & Hackett, T. A. Thalamic connections of the auditory cortex in marmoset monkeys: Core and medial belt regions. J. Comp. Neurol. 496, 72–96 (2006).

28. Lurie, D. I., Pasic, T. R., Hockfield, S. J. & Rubel, E. W. Development of Cat-301 immunoreactivity in auditory brainstem nuclei of the gerbil. J. Comp. Neurol. 380, 319–334 (1997).

29. Stein, J. The visual basis of reading and reading difficulties. Front. Neurosci. 16, 1004027 (2022).

30. Stein, J. & Talcott, J. Impaired neuronal timing in developmental dyslexia - The magnocellular hypothesis. Dyslexia 5, 59–77 (1999).

31. Trussell, L. O. Cellular mechanisms for preservation of timing in central auditory pathways. Curr. Opin. Neurobiol. 7, 487–492 (1997).

32. Winer, J. A. & Morest, D. K. The medial division of the medial geniculate body of the cat: Implications for thalamic organization. J. Neurosci. 3, 2629–2651 (1983).

33. Merigan, W. H. & Maunsell, J. H. R. How parallel are the primate visual pathways? Annu. Rev. Neurosci. 16, 369–402 (1993).

34. Zhang, P., Zhou, H., Wen, W. & He, S. Layer-specific response properties of the human lateral geniculate nucleus and superior colliculus. Neuroimage 111, 159–166 (2015).

35. Livingstone, M. S. & Hubel, D. H. Psychophysical evidence for separate channels for the perception of form, color, movement, and depth. J. Neurosci. 7, 3416–3468 (1987).

36. Arnal, L. H., Flinker, A., Kleinschmidt, A., Giraud, A. L. & Poeppel, D. Human Screams Occupy a Privileged Niche in the Communication Soundscape. Curr. Biol. 25, 2051–2056 (2015).

37. Galaburda, A. M., Menard, M. T. & Rosen, G. D. Evidence for aberrant auditory anatomy in developmental dyslexia. Proc. Natl. Acad. Sci. 91, 8010–8013 (1994).

38. Galaburda, A. & Livingstone, M. Evidence for a Magnocellular Defect in Developmental Dyslexia a. Ann. N. Y. Acad. Sci. 682, 70–82 (1993).

39. Meng, Q. & Schneider, K. A. A specialized channel for encoding auditory transients in the magnocellular division of the human medial geniculate nucleus. Neuroreport 33, 663–668 (2022).

40. Stuart, G. W., McAnally, K. I., McKay, A., Johnston, M. & Castles, A. A test of the magnocellular deficit theory of dyslexia in an adult sample. Cogn. Neuropsychol. 23, 1215–29 (2006).

41. Domínguez-Borràs, J. et al. Human amygdala response to unisensory and multisensory emotion input: No evidence for superadditivity from intracranial recordings. Neuropsychologia 131, 9–24 (2019).

42. Sagaspe, P., Schwartz, S. & Vuilleumier, P. Fear and stop: A role for the amygdala in motor inhibition by emotional signals. Neuroimage 55, 1825–1835 (2011).

43. Rentzsch, J., Jockers-Scherübl, M. C., Boutros, N. N. & Gallinat, J. Test-retest reliability of P50, N100 and P200 auditory sensory gating in healthy subjects. Int. J. Psychophysiol. 67, 81–90 (2008).

44. Phelps, E. A., Ling, S. & Carrasco, M. Emotion facilitates perception and potentiates the perceptual benefits of attention. Psychol. Sci. 17, 292–299 (2006).

45. Lonsdorf, T. B. et al. Don’t fear ‘fear conditioning’: Methodological considerations for the design and analysis of studies on human fear acquisition, extinction, and return of fear. Neurosci. Biobehav. Rev. 77, 247–285 (2017).

46. LeDoux, J. The emotional brain, fear, and the amygdala. Cell. Mol. Neurobiol. 23, 727–738 (2003).

47. Armony, J. L. & Dolan, R. J. Modulation of spatial attention by fear-conditioned stimuli: An event-related fMRI study. Neuropsychologia 40, 817–826 (2002).

48. Pessoa, L. & Adolphs, R. Emotion processing and the amygdala: from a ‘low road’ to ‘many roads’ of evaluating biological significance. Nat. Rev. Neurosci. 11, 773–83 (2010).

49. Malmierca, M. S., Merchán, M. A., Henkel, C. K. & Oliver, D. L. Direct projections from cochlear nuclear complex to auditory thalamus in the rat. J. Neurosci. 22, 10891–10897 (2002).

50. Anderson, L. A., Malmierca, M. S., Wallace, M. N. & Palmer, A. R. Evidence for a direct, short latency projection from the dorsal cochlear nucleus to the auditory thalamus in the guinea pig. Eur. J. Neurosci. 24, 491–498 (2006).

51. Strominger, N. L., Nelson, L. R. & Dougherty, W. J. Second order auditory pathways in the chimpanzee. J. Comp. Neurol. 172, 349–365 (1977).

52. Domínguez-Borràs, J., Rieger, S. W., Corradi-Dell’Acqua, C., Neveu, R. & Vuilleumier, P. Fear spreading across senses: Visual emotional events alter cortical responses to touch, audition, and vision. Cereb. Cortex 27, 68–82 (2017).

53. Pourtois, G., Schettino, A. & Vuilleumier, P. Brain mechanisms for emotional influences on perception and attention: What is magic and what is not. Biol. Psychol. 92, 492–512 (2013).

54. Quirk, G. J., Armony, J. L. & LeDoux, J. E. Fear conditioning enhances different temporal components of tone-evoked spike trains in auditory cortex and lateral amygdala. Neuron 19, 613–624 (1997).

55. Cano, G., Hernan, S. L. & Sved, A. F. Centrally projecting Edinger-Westphal nucleus in the control of sympathetic outflow and energy homeostasis. Brain Sci. 11, (2021).

56. Roelofs, K. Freeze for action: Neurobiological mechanisms in animal and human freezing. Philos. Trans. R. Soc. B Biol. Sci. 372, (2017).

57. Szabadi, E. Modulation of physiological reflexes by pain: role of the locus coeruleus. Front. Integr. Neurosci. 6, 94 (2012).

58. Morris, J. S., Öhman, A. & Dolan, R. J. A subcortical pathway to the right amygdala mediating ‘unseen’ fear. Proc. Natl. Acad. Sci. U. S. A. 96, 1680–1685 (1999).

59. Anderson, L. A. & Linden, J. F. Physiological differences between histologically defined subdivisions in the mouse auditory thalamus. Hear. Res. 274, 48–60 (2011).

60. Garrido, M. I., Barnes, G. R., Sahani, M. & Dolan, R. J. Functional evidence for a dual route to amygdala. Curr. Biol. 22, 129–134 (2012).

61. Kumar, S., von Kriegstein, K., Friston, K. & Griffiths, T. D. Features versus feelings: Dissociable representations of the acoustic features and valence of aversive sounds. J. Neurosci. 32, 14184–14192 (2012).

62. Giraud, A. L. & Poeppel, D. Cortical oscillations and speech processing: Emerging computational principles and operations. Nat. Neurosci. 15, 511–517 (2012).

63. Stein, J. 30612 The Reading Networks and Dyslexia. The Neural Basis of Reading 0 (2010) doi:10.1093/acprof:oso/9780195300369.003.0012.

64. Trevor, C., Arnal, L. H. & Frühholz, S. Terrifying film music mimics alarming acoustic feature of human screams. J. Acoust. Soc. Am. 147, EL540–EL545 (2020).

65. McFadyen, J., Mermillod, M., Mattingley, J. B., Halász, V. & Garrido, M. I. A rapid subcortical amygdala route for faces irrespective of spatial frequency and emotion. J. Neurosci. 37, 3864–3874 (2017).

66. Bayle, D. J., Schoendorff, B., Hénaff, M. A. & Krolak-Salmon, P. Emotional facial expression detection in the peripheral visual field. PLoS One 6, (2011).

67. Bayle, D. J., Henaff, M. A. & Krolak-Salmon, P. Unconsciously perceived fear in peripheral vision alerts the limbic system: A MEG study. PLoS One 4, (2009).

68. Kragel, P. A. et al. A human colliculus-pulvinar-amygdala pathway encodes negative emotion. Neuron 109, 2404–2412.e5 (2021).

69. Khalil, V. et al. Subcortico-amygdala pathway processes innate and learned threats. Elife 12, 1–26 (2023).

70. Li, W. & Keil, A. Sensing fear: fast and precise threat evaluation in human sensory cortex. Trends Cogn. Sci. 27, 341–352 (2023).

71. Carretié, L. et al. Fast unconscious processing of emotional stimuli in early stages of the visual cortex. Cereb. Cortex 32, 4331–4344 (2022).

72. Weinberger, N. M. The medial geniculate, not the amygdala, as the root of auditory fear conditioning. Hear. Res. 274, 61–74 (2011).

73. Domínguez-Borràs, J., Bourgeois, A. & Vuilleumier, P. Affective Biases in Attention, Cognitive Control, and Awareness. in The Cambridge Handbook of Human Affective Neuroscience 385–407 (Cambridge University Press, 2025). doi:10.1017/9781009342919.025.

74. Hockfield, S. & Sur, M. Monoclonal antibody cat-301 identifies Y-cells in the dorsal lateral geniculate nucleus of the cat. J. Comp. Neurol. 300, 320–330 (1990).

75. Shen, L., Liu, D. & Huang, Y. Hypothesis of subcortical visual pathway impairment in schizophrenia. Med. Hypotheses 156, (2021).

76. Javitt, D. C. & Freedman, R. Sensory processing dysfunction in the personal experience and neuronal machinery of schizophrenia. Am. J. Psychiatry 172, 17–31 (2015).

77. Fecteau, S., Belin, P., Joanette, Y. & Armony, J. L. Amygdala responses to nonlinguistic emotional vocalizations. Neuroimage 36, 480–487 (2007).

78. Pannese, A., Grandjean, D. & Frühholz, S. Subcortical processing in auditory communication. Hear. Res. 328, 67–77 (2015).

79. Sauter, D. A., Eisner, F., Calder, A. J. & Scott, S. K. Perceptual cues in nonverbal vocal expressions of emotion. Q. J. Exp. Psychol. 63, 2251–2272 (2010).

80. Belin, P., Fillion-Bilodeau, S. & Gosselin, F. The Montreal Affective Voices: A validated set of nonverbal affect bursts for research on auditory affective processing. Behav. Res. Methods 40, 531–539 (2008).

81. Büchel, C., Morris, J., Dolan, R. J. & Friston, K. J. Brain systems mediating aversive conditioning: An event-related fMRI study. Neuron 20, 947–957 (1998).

82. Groppe, D. M., Urbach, T. P. & Kutas, M. Mass univariate analysis of event-related brain potentials/fields I: a critical tutorial review. Psychophysiology 48, 1711–1725 (2011).

83. Maris, E. & Oostenveld, R. Nonparametric statistical testing of EEG- and MEG-data. J. Neurosci. Methods 164, 177–190 (2007).

84. Oostenveld, R., Fries, P., Maris, E. & Schoffelen, J. M. FieldTrip: Open source software for advanced analysis of MEG, EEG, and invasive electrophysiological data. Comput. Intell. Neurosci. 2011, (2011).

85. Costa-Faidella, J., Baldeweg, T., Grimm, S. & Escera, C. Interactions between ‘what’ and ‘when’ in the auditory system: Temporal predictability enhances repetition suppression. J. Neurosci. 31, 18590–18597 (2011).

86. Bates, D., Mächler, M., Bolker, B. M. & Walker, S. C. Fitting linear mixed-effects models using lme4. J. Stat. Softw. 67, (2015).

87. Schielzeth, H. et al. Robustness of linear mixed-effects models to violations of distributional assumptions. Methods Ecol. Evol. 11, 1141–1152 (2020).

88. Paraskevoudi, N. & SanMiguel, I. Sensory suppression and increased neuromodulation during actions disrupt memory encoding of unpredictable self-initiated stimuli. Psychophysiology 60, 1–25 (2023).

89. Binda, P., Pereverzeva, M. & Murray, S. O. Attention to bright surfaces enhances the pupillary light reflex. J. Neurosci. 33, 2199–2204 (2013).

90. Knapen, T. et al. Cognitive and ocular factors jointly determine pupil responses under equiluminance. PLoS One 11, 1–13 (2016).

