## Supplementary Information for "An auditory “low road” for threat in humans sensitive to fast temporal cues"

**This PDF file includes:**

Supplementary Information text
Supplementary Figures 1 to 3
Supplementary Tables 1 to 3
Supplementary References

**Supplementary Information Text**

**Supplementary Results**

***Study 1. No influence of intrinsic physical properties of highAM and lowAM on the observed ERP effects***

There were no differences in early neural responses between highAM and lowAM during the Pre-Conditioning phase, where no emotional (conditioned) stimuli were present. No main or interaction effect was found (main effect of AM: $p$ = .214, $r^2$ < .01; main effect of CS: $p$ = .702, $r^2$ < .01; AM x CS: $p$ = .421, $r^2$ < .01) between the four different conditions during Pre-Conditioning, suggesting that no effect driven by their intrinsic physical properties was present in the early neural response.

**Supplementary Materials and Methods**

***Study 1. Identifying fast threat responses to highAM cues (EEG and pupillometry)***

**Details on EEG preprocessing**

EEG activity was recorded at a sampling rate of 500 Hz in a Neuroscan SynAmps RT system (NeuroScan, Compumedics, Charlotte, NC, USA), from 64 Ag/AgCl electrodes placed in a nylon cap following the extended 10-20 system. The horizontal electrooculogram (EOG) was measured with two electrodes placed on the outer canthi of the eyes, and the vertical EOG with two electrodes placed on above and below the left eye, all referenced to the common reference. One electrode placed at the tip of the nose served as online reference and the AFz electrode was used as the ground. All impedances were kept below 10 kΩ.

Preprocessing was done in MATLAB using EEGLab[1]. Signal was bandpass filtered at 0.1-100 Hz for long latency responses (FIR orders 9056 and 1812, respectively) and at 10-200 Hz for middle latency responses (FIR order 1812), all using a Kaiser window (β=5.653). Based on the 64 electrodes, independent component analysis was used to remove eye movements and muscle activity, using the Second Order Blind Identification (SOBI), after visual inspection. Epochs were defined from − 200 ms to 1000 ms relative to stimulus (CS) onset, and baseline-corrected (200 ms prior to the sound onset). Trials containing the WN (condition CS+) were not included in the analysis. Malfunctioning electrodes were interpolated with spherical interpolation. Epochs exceeding ± 75 μV in more than 60% of the electrodes were discarded. Event-related potentials (ERPs) were obtained by averaging the remaining epochs within each participant for each condition (for trials with correct responses). Average trial counts per condition were 53 ± 9.4 for long-latency responses and 63 ± 3.8 for the middle-latency responses (Supplementary Table 2).

**Number of trials per condition**

The Pre-Conditioning phase consisted of 208 trials (52 trials per condition, with 26 trials containing left-lateralized voices, and 26 right-lateralized voices), distributed in four blocks. The Conditioning phase comprised 416 trials across nine blocks. This phase included additional CS+ alone trials (52 highAM, 52 lowAM), and CS+ trials followed by the US (52 CS+ highAM, 52 CS+ lowAM). Because the Conditioning phase included more CS+ than CS- trials (due to the addition of CS+ pairings with the US), we increased the number of CS- trials to minimize expectancy or oddball effects arising from this imbalance [2]. Specifically, the Conditioning phase included 104 CS- highAM and 104 CS- lowAM trials. We further confirmed that any differences in trial numbers across conditions did not bias EEG results by including number of trials as covariate ($p$ > .05). The additional Extinction phase consisted of 40 trials (5 trials per condition, one block).

### Assessing the potential influence of intrinsic physical properties of highAM versus lowAM stimuli

To check for possible differences in early neural responses between highAM and lowAM driven by their intrinsic physical properties, we performed a LMM into the four different sounds during Pre-Conditioning phase. Each model included CS and AM as fixed effects, with Participant modeled as a random effect.

### *Study 2. Associating an MGB-BLA pathway with threat-related responses to highAM cues (fMRI and diffusion tractography)*

#### Subjects

Thirty-six healthy volunteers were mostly recruited among University of Barcelona students. All were right-handed, with normal hearing and no previous neurological or psychiatric disorders. Six participants were excluded from the analysis due to low signal-to-noise ratio in the MRI data and drowsiness during task performance. Thus, the final sample consisted of thirty-one (18 females; age range 18-39, mean age = 24.35). Informed consent was obtained, and all procedures were approved by the ethics committee of the University of Barcelona, in accordance with the Declaration of Helsinki (2024). All participants received monetary compensation for their participation.

#### Stimuli

One 400-ms binaural excerpt of female non-verbal vocal utterance with neutral prosody, extracted from the Montreal Affective Voices database[3], served as conditioned stimuli (CS). It was digitally modulated in amplitude at high (40 Hz) and low (10 Hz) amplitude modulation (AM) rates. Stimuli were calibrated for each participant to be well noticeable in the scanner (mean: 20 dB SPL above hearing threshold). Additionally, a 400-ms binaural burst of an unpleasant, but not painful, loud white noise and a pool of different negative images from the International affective picture system (IAPS; Bradley & Lang, 2017[4]) served as unconditioned stimulus (US). Images served to reinforce the conditioning effect inside the scanner despite background noise.

#### Procedure and task

Participants were instructed to respond, as quickly and accurately as possible, on which side they heard a voice (left or right, by pressing a keyboard button with their right index or middle finger, respectively), while maintaining their gaze on a permanent fixation cross at the center of the screen. Voices were presented in trials of 5000±800-ms duration.

The task consisted of a fear conditioning paradigm with a Pre-Conditioning and a Conditioning phase. During Conditioning, participants were presented with both highAM and lowAM voices, and these were either paired (CS+) or unpaired (CS-) with the unpleasant loud white noise and one IAPS aversive image (US). Contrarily to Study 1, stimulus assignment as either CS+ or CS- was based on the source location of each voice instead of the gender of the stimulus voice, and similarly counterbalanced across participants (i.e., 50% were presented with highAM and lowAM CS+ left-sided voices, and with highAM and lowAM CS- right-sided voices). The CS-US pairing conferred aversive emotional value to the CS+, whereas the CS- remained neutral. During Pre-Conditioning, the same CS stimuli were presented without the US. The fear conditioning was conducted with a 50% partial reinforcement.

The Pre-Conditioning phase consisted of 104 trials (26 trials per condition), distributed into 3 blocks. The Conditioning phase consisted of 104 trials distributed across 3 blocks. Similarly, as in Study 1, this phase consisted of more trials than in Pre-Conditioning given the presence of additional CS+ alone voices (resulting into 52 CS+ alone highAM, 52 CS+ alone lowAM, but also trials containing the US; 52 CS+ highAM, and 52 CS+ lowAM trials).

Voices were presented in pseudo-random order, ensuring no more than two consecutive repetitions of the same stimulus to prevent habituation[5]. Voices were delivered binaurally, but with manipulated amplitude-difference so they were perceived as lateralized. In turn, if participants did not respond within 2 seconds after voice onset, response was counted as a miss.

At the end of each block of the Conditioning phase, participants rated, on a scale from 1 (never) to 5 (always), how likely each of the voices used in the experiment was followed by the US, so as to measure their awareness of CS-US contingency. Finally, at the beginning and at the end of the Conditioning phase, participants were asked to rate all voices for valence and arousal on a scale from 1 (very positive – little stimulating) to 5 (very negative – very stimulating).

**Data acquisition**

Whole-brain MRI data were acquired on a 3T Philips Ingenia CX scanner equipped with a 32-channel whole-head coil, located at the BarcelonaBeta brain research center (https://www.barcelonabeta.org/es). The imaging protocol included high-resolution structural T1-weighted (T1w) scans, functional MRI (fMRI), and diffusion-weighted imaging (DWI) sequences. The T1-weighted structural images were obtained using a Turbo Field Echo (TFE) sequence with the following parameters: repetition time (TR) = 9.9 ms, echo time (TE) = 4.6 ms, flip angle (FA) = 8°, field of view (FoV) = 240 mm, voxel size = 0.9 mm isotropic, and 200 slices. The functional MRI scans were acquired with TR = 2660 ms, TE = 33 ms, FA = 80°, FoV = 240 mm, voxel size = 1.8 mm isotropic, and 66 slices with a 0.36 mm inter-slice gap. A multiband acceleration factor of 2 was applied to shorten acquisition time. For the diffusion-weighted imaging, a total of 108 volumes were acquired in the anterior-to-posterior phase-encoding direction. This included 96 diffusion-weighted directions distributed across multiple b-values: 8 directions at b = 500 s/mm², 8 at b = 1000 s/mm², 16 at b = 2000 s/mm², and 64 at b = 3000 s/mm². Additionally, 12 non-diffusion-weighted (b = 0 s/mm²) images were included in the main acquisition. For distortion and motion correction, two extra b = 0 s/mm² images were acquired with opposite phase-encoding directions (anterior-to-posterior and posterior-to-anterior). DWI acquisition parameters for both the diffusion-weighted and b0 images were as follows: TR = 7463 ms, TE = 89 ms, FA = 90°, voxel size = 1.65 mm isotropic, 81 slices with no gap, and a multiband acceleration factor of 3.

**Diffusion MRI preprocessing**

Preprocessing of diffusion-weighted images was performed using functions from the MRtrix3 software. First, denoising was conducted using the dwidenoise function, which applies patch-based principal component analysis grounded in random matrix theory to exploit data redundancy[6,7]. Rician background noise was subsequently removed using mrcalc. Gibbs ringing artifacts were corrected with mrdegibbs[8]. Susceptibility-induced distortions and motion artifacts were corrected using FSL's topup and eddy tools[9], integrated via the dwifslpreproc function. B1 field inhomogeneity correction was performed using dwibiascorrect, and a brain mask was generated with dwi2mask to constrain subsequent analysis to brain tissue. Finally, the T1w anatomical image was rigidly aligned to the DWI data using ANTs.

We estimated response functions for grey matter, white matter and cerebrospinal fluid (CSF) using the multi-shell multi-tissue constrained spherical deconvolution (MSMT-CSD) algorithm for each participant[10]. Using these response functions, we then produced individual multi-tissue fiber orientation distribution functions (fODFs) with the MSMT-CSD algorithm, which represent possible fiber directions with corresponding weights in each voxel. Finally, we performed joint bias field correction and global intensity normalization of the multi-tissue compartment parameters using the mtnormalise command in MRtrix3.

**Region of interest (ROI) definition**

ROI definition involved processing each subject's T1w aligned anatomical image to obtain the ROIs for tractography. This process takes as input the subject's T1w image and predefined ROIs in MNI space, and outputs a segmented T1w image along with the ROIs in individual subject T1w space. Because the T1w image was already rigidly aligned to the DWI data, the resulting ROIs were likewise aligned and suitable for use in tractography analyses.

Cortical and subcortical segmentation of the T1-weighted (T1w) anatomical images was performed using FreeSurfer (http://surfer.nmr.mgh.harvard.edu/) The medial geniculate nucleus (MGB) ROI was extracted using FreeSurfer's thalamic segmentation module, which utilizes a probabilistic atlas derived from histological data and high-resolution ex vivo MRI[11]. The basolateral amygdala (BLA) ROI was defined using the amygdala segmentation developed by Saygin et al.[12], also implemented in FreeSurfer. The lateral, basal, and accessory basal subregions were combined to form a single BLA ROI. To improve the neuroanatomical specificity of tract reconstruction, exclusion ROIs were applied based on the anatomical location and expected connectivity of the MGB-BLA pathway. These exclusion criteria were informed by initial visual inspection and are detailed in Supplementary Table 3.

**Diffusion MRI analysis**

Diffusion MRI tractography was conducted using MRtrix3, leveraging both the preprocessed DWI data and anatomically defined ROIs to reconstruct the MGB–BLA pathway bilaterally. Whole-brain tractograms were first generated using the iFOD2 probabilistic tractography algorithm, seeding 10 million streamlines with a fiber orientation distribution (FOD) amplitude cutoff of 0.06. To reduce the incidence of false positive and prematurely terminated streamlines, we employed the Anatomically Constrained Tractography (ACT) framework using a five-tissue-type (5TT) segmentation of the T1-weighted image (cortical and subcortical gray matter, white matter, CSF, and pathological tissue; Smith et al.[13]). In the same line, we produced ROI to ROI tractograms for each participant planting a seed point 25.000 times in each voxel of the seeding ROI (MGB) with the following parameters: step size 0.3 mm, maximum fiber length 80 mm, minimum fiber length 5 mm, FOD amplitude threshold of 0.1, and angle threshold of 45 degrees. Resulting tractograms were constrained using inclusion and exclusion ROIs (see Supplementary Table 3), ensuring biologically plausible reconstructions of the MGB-BLA pathway. To quantify structural connectivity, the targeted tractograms were combined with the whole-brain tractogram, and the SIFT2 algorithm was applied. This method assigns a weight to each streamline to ensure quantitative consistency with the underlying fiber density[14]. Streamline weights were finally multiplied by the global scaling factor μ, computed from the whole-brain tractogram, to convert them into quantitatively meaningful measures of fiber bundle capacity (FBC; referred to as fiber density) that are comparable across subjects.

**Functional MRI preprocessing**

We used statistical parametric mapping (SPM) software (version 12; Welcome Department of Cognitive Neurology, London, United Kingdom) for preprocessing and statistical analysis of functional images from our complementary study. Volumes were slice-time–corrected to the first data point and realigned using the "register to mean" option. Anatomical images were coregistered to the functional image by applying a normalized mutual information three-dimensional rigid-body transformation. Brain segmentation and posterior normalization to the Montreal Neurological Institute (MNI) space was performed to the functional images resampled to a 2-mm³ voxel size and spatially smoothed with a 6 mm³ full width at half-maximum (FWHM) Gaussian kernel.

The interleaved sequence used to acquire functional time series made it a prerequisite to use slice-time correction as a first preprocessing step[15]. Slice-timing correction methods can successfully compensate for slice-timing effects[16].

**Functional MRI analysis**

For individual analysis (first level), trial onsets from the 8 conditions of interest (see Supplementary Table 2) and the 2 conditions in which the US was presented were modeled in the design matrix as separate regressors. To account for movement-related variance, the first-level GLM also included realignment parameters from each session [x, y, and z translations and pitch, roll, and yaw rotations] and the temporal modulator as covariate of interest.

After model estimation, contrast images were calculated for each experimental condition (versus baseline). The resulting individual contrast images were then entered into a flexible factorial design at the second level. Fiber density values of the left and right MGB – BLA tract, previously extracted for each participant (see above) were included as two continuous covariates, and modeled with factor-specific interaction terms (fiber density × factor AM, with the levels highAM and lowAM and fiber density x factor CS, with levels CS+ and CS-), again computed as the difference between Conditioning and Pre-Conditioning. We tested whether the strength of the association between fiber density and hemodynamic response in amygdala differed across AM rate (highAM versus lowAM) and CS type (CS+ versus CS-) with special focus on its interaction ([highAM CS+ > highAM CS-] – [lowAM CS+ < lowAM CS-]). To do so, the model included interaction terms between each fiber density and each experimental factor, and contrasts compared the corresponding interaction regressors. To assess for threat-related responses, we focused particularly on amygdala activity[17,18]. To this end, small volume correction (SVC) was applied with the anatomical mask of bilateral or right amygdala extracted from the WFU pickatlas (RRID: SCR_007378) when appropriate. Peak-level significance mask was assessed using family-wise error (FWE) correction for multiple comparisons, with a statistical threshold of $p < 0.05$ FWE-SVC as significance criterion. A whole analysis (FWE-corrected) was also computed for the contrasts of interest.

### Supplementary Figures

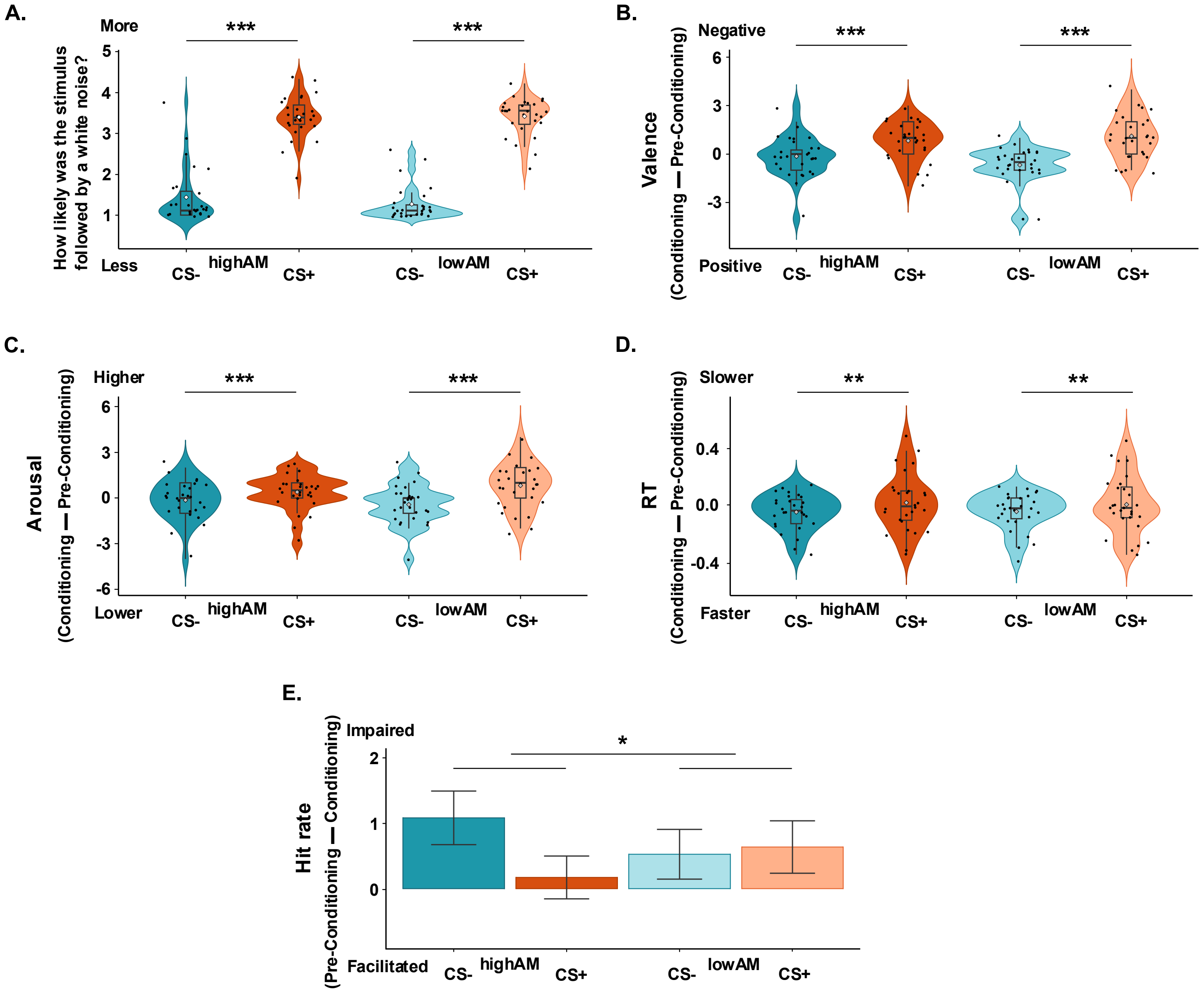


**Supplementary Figure 1. Study 1 behavioral results.** Contingency (A), valence (B) and arousal (C) ratings. (D) Response times. (E) Hit rates. (A – D) The y axis shows the difference Conditioning – Pre-Conditioning. (E) The y axis shows the opposite difference (Pre-Conditioning – Conditioning) for a better visualization of the effect. (* $p \leq .05$, ** $p \leq .01$, *** $p \leq .001$).

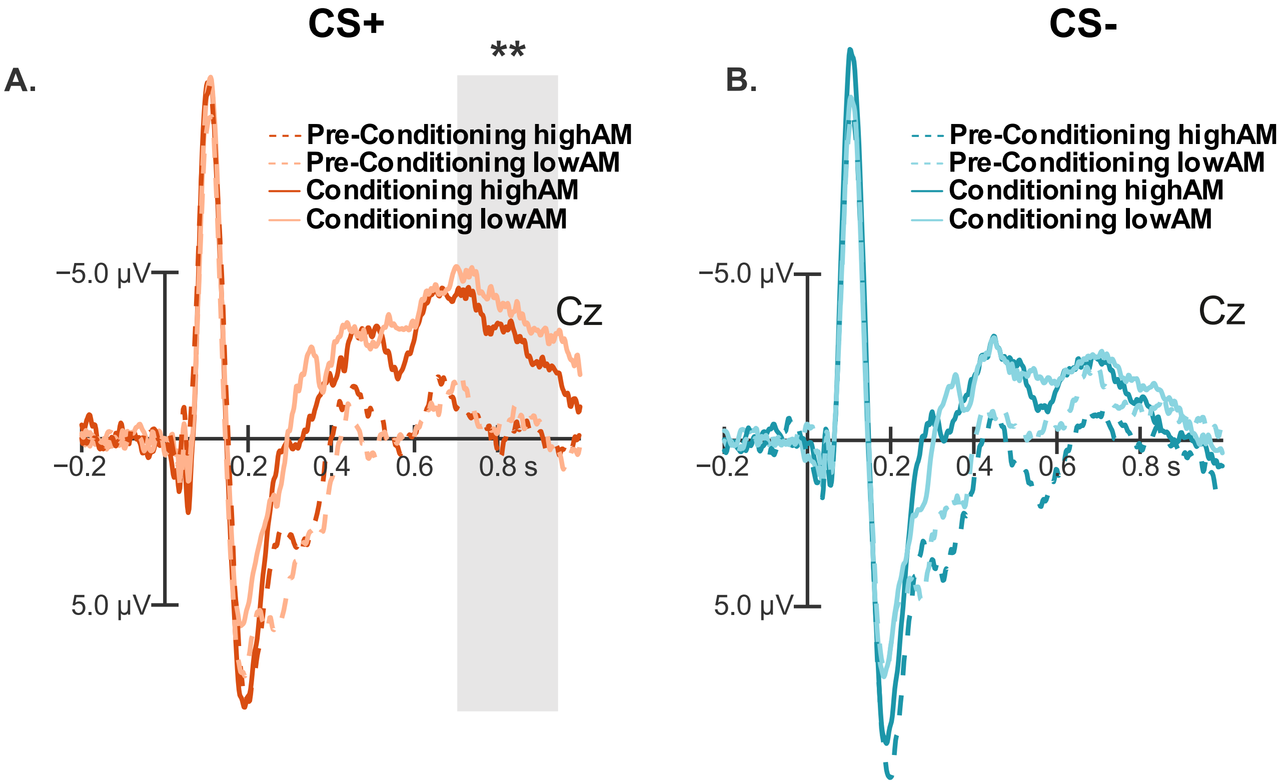


**Supplementary Figure 2. Study 1 electrophysiological data.** (A) Neural responses for emotional (CS+) stimuli. (B) Neural responses for emotional (CS-) stimuli. Grey areas indicate significant effects (** $p \leq .01$).

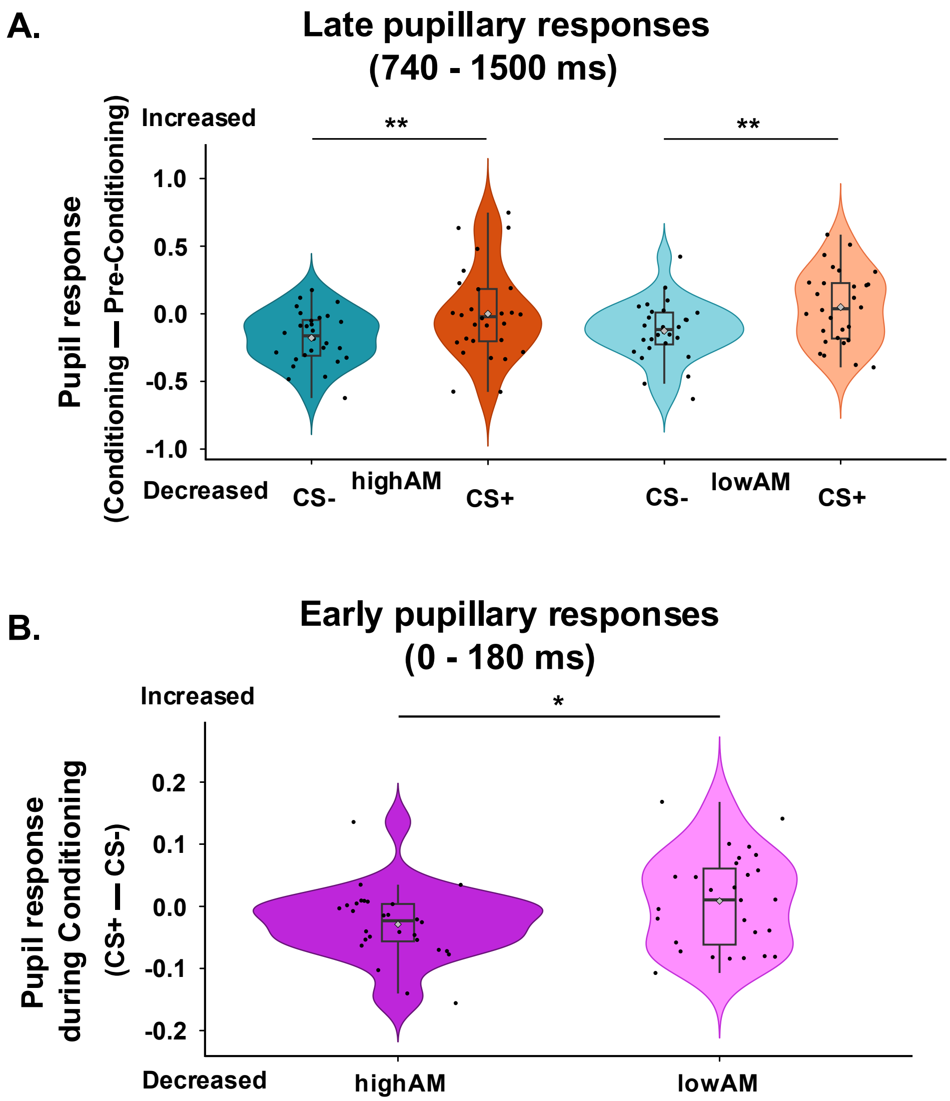


**Supplementary Figure 3. Study 1 pupillary results.** (A) Late pupillary responses (740 - 1500 ms post-stimulus average response). The y axis shows the difference Conditioning – Preconditioning. (B) Early pupillary responses (0 – 180 ms post-stimulus average response). The y axis shows the difference CS+ – CS- during the Conditioning phase. (* $p \leq .05$, ** $p \leq .01$).

**Supplementary Table 1.** Effects of MGB – BLA covariate in amygdala response

| Side | Area | x | y | z | T value | Size (voxels) | P value |
|---|---|---|---|---|---|---|---|
| rMGB – BLA x Interaction AM x CS ([highAM CS+ > highAM CS-] – [lowAM CS+ < lowAM CS-]) | | | | | | | |
| R | amygdala | 28 | -6 | -22 | 4.59 | 80 | < .001 |
| rMGB – BLA x reversed Interaction AM x CS ([highAM CS+ < highAM CS-] – [lowAM CS+ > lowAM CS-]) | | | | | | | |
| | | | | | | | ns |
| rMGB – BLA x highAM (highAM CS+ > highAM CS-) | | | | | | | |
| R | amygdala | 26 | -6 | -20 | 4.23 | 87 | .002 |
| rMGB – BLA x lowAM (lowAM CS+ > lowAM CS-) | | | | | | | |
| | | | | | | | ns |

Note: All coordinates reported in MNI space and $p < 0.05$ corrected for small volume based on amygdala masks from WFU pickatlas (see SI Materials Methods section). L = left, R = right, ns = non-significant.

**Supplementary Table 2.** Study 1: number of trials accounted per condition for the ERP analyses

| | P50 analysis | | | | Full ERP analysis | | | |
|---|---|---|---|---|---|---|---|---|
| Condition | Mean | SD | Min | Max | Mean | SD | Min | Max |
| **highAM CS+ Pre** | 50.43 | 2.73 | 40 | 52 | 43.93 | 6.87 | 30 | 52 |
| **highAM CS- Pre** | 50.32 | 2.58 | 43 | 52 | 43.25 | 6.61 | 27 | 50 |
| **lowAM CS+ Pre** | 50.21 | 2.79 | 40 | 52 | 43.29 | 7,03 | 25 | 52 |
| **lowAM CS- Pre** | 49.86 | 2.99 | 40 | 52 | 44.46 | 5.72 | 32 | 52 |
| **highAM CS+ Cond** | 48.93 | 3.65 | 38 | 52 | 41.61 | 8.25 | 23 | 52 |
| **highAM CS- Cond** | 98.79 | 5.84 | 83 | 104 | 81.43 | 16.12 | 48 | 104 |
| **lowAM CS+ Cond** | 48.75 | 3.54 | 39 | 52 | 42.21 | 8.37 | 21 | 52 |
| **lowAM CS- Cond** | 98.32 | 6.08 | 82 | 104 | 83.46 | 16.49 | 47 | 103 |

Abbreviations: Pre = Pre-Conditioning phase, Cond = Conditioning phase, SD = Standard deviation, Min = Minimum, Max = Maximum

**Supplementary Table 3.** Study 2: tracts and ROIs included to identify them.

| Tract name | Inclusion ROI 1 | Inclusion ROI 2 | Exclusion ROIs |
|---|---|---|---|
| MGB – BLA | ***MGB*** | **BLA** | PuA, PuM, PuL, PuI, MD, AV, VA, LD, VPL |

Abbreviations: PuA = anterior pulvinar, PuM = medial pulvinar, PuL = lateral pulvinar, PuI = inferior pulvinar, MD = mediodorsal nucleus, AV = anteroventral nucleus, VA = ventral anterior nucleus, LD = laterodorsal nucleus, VPL = ventral posterolateral nucleus

**Supplementary References**


1. Delorme, A. & Makeig, S. EEGLAB: An open source toolbox for analysis of single-trial EEG dynamics including independent component analysis. *J. Neurosci. Methods* **134**, 9–21 (2004).
2. Näätänen, R., Kujala, T. & Winkler, I. Auditory processing that leads to conscious perception: a unique window to central auditory processing opened by the mismatch negativity and related responses. *Psychophysiology* **48**, 4–22 (2011).
3. Belin, P., Fillion-Bilodeau, S. & Gosselin, F. The Montreal Affective Voices: A validated set of nonverbal affect bursts for research on auditory affective processing. *Behav. Res. Methods* **40**, 531–539 (2008).
4. Bradley, M. M. & Lang, P. J. International Affective Picture System BT - Encyclopedia of Personality and Individual Differences. in (eds. Zeigler-Hill, V. & Shackelford, T. K.) 1–4 (Springer International Publishing, 2017). doi:10.1007/978-3-319-28099-8_42-1.
5. Lonsdorf, T. B. *et al.* Don't fear 'fear conditioning': Methodological considerations for the design and analysis of studies on human fear acquisition, extinction, and return of fear. *Neurosci. Biobehav. Rev.* **77**, 247–285 (2017).
6. Cordero-Grande, L., Christiaens, D., Hutter, J., Price, A. N. & Hajnal, J. V. Complex diffusion-weighted image estimation via matrix recovery under general noise models. *Neuroimage* **200**, 391–404 (2019).
7. Veraart, J. *et al.* Denoising of diffusion MRI using random matrix theory. *Neuroimage* **142**, 394–406 (2016).
8. Kellner, E., Dhital, B., Kiselev, V. G. & Reisert, M. Gibbs-ringing artifact removal based on local subvoxel-shifts. *Magn. Reson. Med.* **76**, 1574–1581 (2016).
9. Smith, S. M. *et al.* Advances in functional and structural MR image analysis and implementation as FSL. *Neuroimage* **23**, S208–S219 (2004).
10. Jeurissen, B., Tournier, J. D., Dhollander, T., Connelly, A. & Sijbers, J. Multi-tissue constrained spherical deconvolution for improved analysis of multi-shell diffusion MRI data. *Neuroimage* **103**, 411–426 (2014).
11. Iglesias, J. E. *et al.* A probabilistic atlas of the human thalamic nuclei combining ex vivo MRI and histology. *Neuroimage* **183**, 314–326 (2018).
12. Saygin, Z. M. *et al.* High-resolution magnetic resonance imaging reveals nuclei of the human amygdala: manual segmentation to automatic atlas. *Neuroimage* **155**, 370–382 (2017).
13. Smith, R. E., Tournier, J.-D., Calamante, F. & Connelly, A. Anatomically-constrained tractography: improved diffusion MRI streamlines tractography through effective use of anatomical information. *Neuroimage* **62**, 1924–38 (2012).
14. Smith, R. E., Tournier, J.-D., Calamante, F. & Connelly, A. SIFT2: Enabling dense quantitative assessment of brain white matter connectivity using streamlines tractography. *Neuroimage* **119**, 338–51 (2015).
15. Parker, D. B. & Razlighi, Q. R. The Benefit of Slice Timing Correction in Common fMRI Preprocessing Pipelines. *Front. Neurosci.* **13**, 821 (2019).
16. Sladky, R. *et al.* Slice-timing effects and their correction in functional MRI. *Neuroimage* **58**, 588–594 (2011).
17. LeDoux, J. E., Farb, C. & Ruggiero, D. A. Topographic organization of neurons in the acoustic thalamus that project to the amygdala. *J. Neurosci.* **10**, 1043–1054 (1990).
18. Ledoux, J. E. & Brown, R. A higher-order theory of emotional consciousness. *Proc. Natl. Acad. Sci. U. S. A.* **114**, E2016–E2025 (2017).
